# Reading about sensations recruits the posterior insula: An intracranial EEG study

**DOI:** 10.1101/2025.03.04.641371

**Authors:** W. Dupont, V. Dornier, R. Palluel-Germain, A. Robin, J-P. Lachaux, P. Kahane, L. Minotti, JR. Vidal, M. Perrone-Bertolotti

## Abstract

When we read, how do words and sentences become concepts? Contrary to theoretical perspectives that conceptualize meaning and concepts as symbolic and abstract, embodied approaches propose that semantic access relies on mental simulations of bodily experiences□associated with the word’s meaning (e.g., simulating the act of running when we read the word ‘run’ or feeling pain when reading the word ‘burn’). If this view of cognition underpins conceptual processing, then sensory and motor cortices should play an early role during language comprehension. To test this idea, we recorded intracranial neural activity (High-Frequency Activity, 50–150 Hz) in epileptic patients while they read short sentences from different semantic categories: abstract-, action- and sensation-related. We hypothesized that reading sensation-related sentences (e.g., “he burns me”) would preferentially involve the posterior insula — a key region for processing affective and painful sensations. Our results, revealed an early functional dissociation between the anterior and posterior insula: the anterior insula showed a non-selective response across all semantic categories, whereas the posterior insula presented an early selective response (around 170 ms) for the sensation-related ones. Furthermore, the intensity of this selective neural response in the posterior insula correlated with the intensity of subjective perceived pain. These results provide direct intracranial evidence suggesting that language is deeply rooted in body-environment interactions.

## Introduction

Contrary to theoretical perspectives that conceptualize meaning and concepts as symbolic and abstract^1^, embodied approaches propose that semantic access relies on mental simulations of bodily experiences associated with the word’s meaning^2,3^ (e.g., simulating the act of running when we read the word ‘run’ or feeling pain when reading the word ‘burn’). If this view of cognition underpins conceptual processing, sensory and motor cortices should play an early role in language comprehension. In this line, several investigations have provided indirect evidence that action word comprehension modulates the motor pathway excitability in Transcranial Magnetic Stimulation studies^4–9^, and the involvement of motor cortex in neuro-imaging studies^10–14^. Additionally, using fMRI, researchers have demonstrated that processing pain-related words is associated with neural activity changes within sensory cortices^15–17^.

To test this idea, we recorded direct intracranial neural activity (High-Frequency Activity -HFA- [50–150 Hz]) in epileptic patients while they read short sentences from different semantic categories: abstract-, action- and sensation-related. Specifically, we analyzed insular cortices activity — a brain region involved in processing bodily and painful sensations^18–22^ — while patients read sensory-related sentences (e.g., “*He burns me*”). We hypothesized that if language comprehension relies on mental simulations grounded in bodily experiences, then reading about sensory experiences (e.g., “he burns me”) should activate brain cortices largely involved in actual bodily sensation, such as the insular cortex. More specifically, we expected that reading sensation-related sentences would preferentially involve an early posterior insula HFA.

## Results and discussion

Sixteen patients (see Table 1 in STAR Methods for demographic details) with drug-resistant epilepsy took part in a Semantic Categorization Task. During the task, participants read minimal sentences including a pronoun-verb segment while brain activity was recorded from a total of 3,329 electrode contacts (intracranial electroencephalography - iEEG) widely distributed across the cortex. Sentences pertained to three semantic categories: abstract (e.g., ‘I hope’), action (e.g., ‘I grasp’), and sensation (e.g., ‘I suffer’). We analyzed High-Frequency Activity (HFA, 50-150 Hz) – as a proxy of population-level spiking activity and a marker of cognitive processing^23^ – to investigate semantic category-specific brain responses. We focused our analyses on the anterior and posterior insula, which are key regions in interoception and pain processing^18,19^. We also evaluated HFA selectivity in functional control regions including: the Inferior Frontal gyrus in the *pars* triangularis for language processing^24,25^, the ventral occipito-temporal cortex for visual word recognition^26–28^, and the precentral gyrus (motor cortex) for action-related processing^29,30^. Experimental procedure and electrode contacts distribution within the insula are illustrated in Figure 1 (the total electrodes distribution is available in supplemental materials, see Figure S1). Behavioral performances were assessed by sentence categorization accuracy (±SEM), which was 83.53% ± 4.23% for abstract, 84.02% ± 3.72% for action, and 75.73% ± 3.50% for sensation-related sentences, indicating successful task execution (see Table S1 for details).

**Figure 1.**
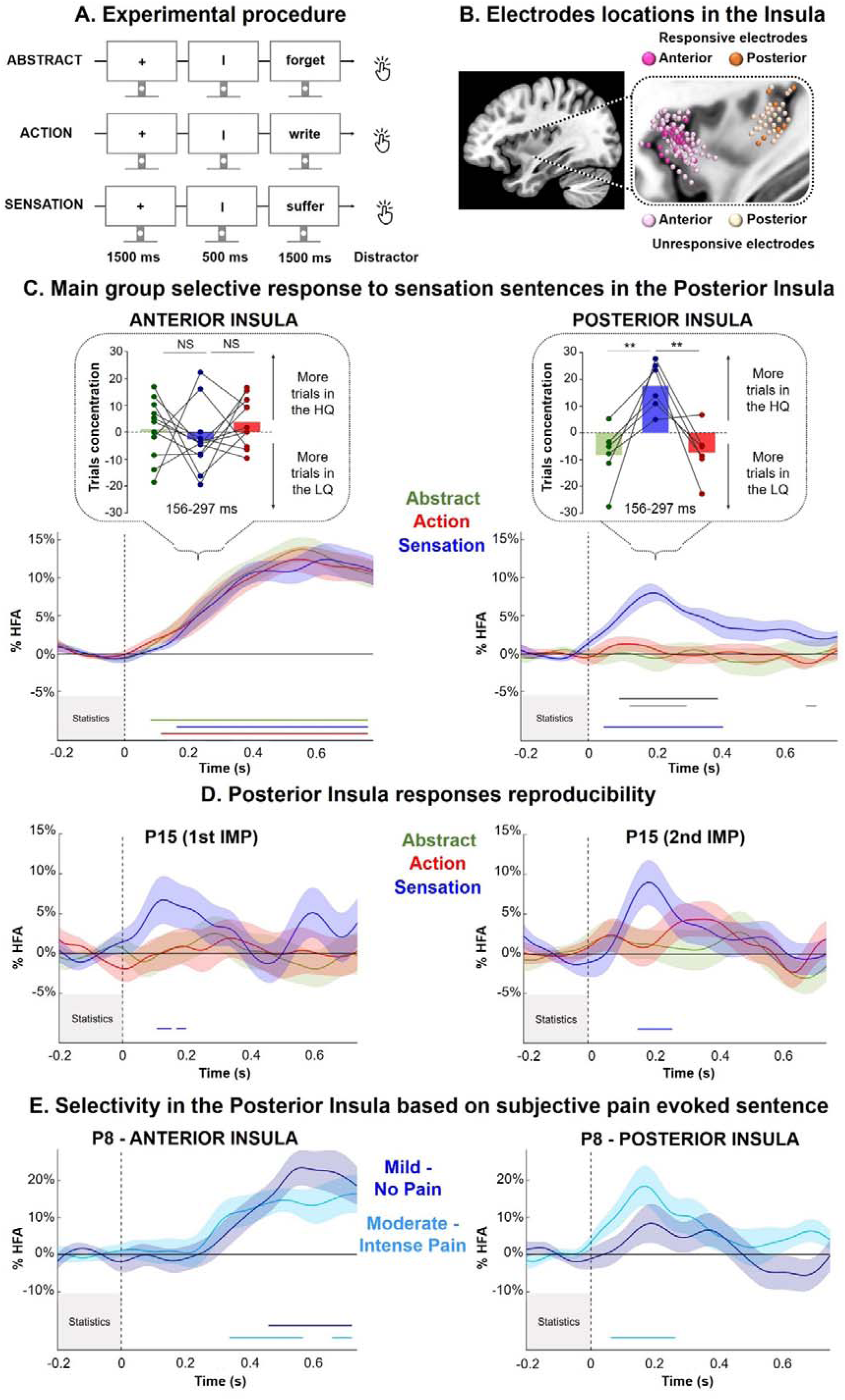
(A) Experimental procedure for the behavioral semantic task, featuring examples of action, abstract and sensation-related sentences. (B) Electrode contact localizations in the anterior and posterior insula, plotted on a standard MRI brain. Electrode contacts in the anterior and posterior insula are colored pink and orange, respectively. Dark and light shades indicate responsives and unresponsives electrode contacts, respectively. (C) Selective neural response for the sensation-related sentences in the anterior and the posterior insula from 6 averaged recordings. For each site, the plots show HFA [50–150 Hz] modulation averaged across trials for Abstract (green), Action (red) and Sensation (bleu) sentences. Amplitude is expressed as a percentage relative to the average amplitude measured across the entire recording session, with shaded areas representing the Standard Error of the Mean (SEM). Colored lines below the HFA plots indicate statistically significant difference between the semantic sentence categories and the baseline, whereas the black and grey lines indicate significant difference between semantic categories. (D) Reproducibility of response selectivity in the posterior insula across two sessions in a single patient (three months separated without resection). (E) Response graduation in the posterior insula as a function of subjective pain evoked by sensation-related sentences. Sensation sentences are subjectively rated by participants as describing very intense pain (light blue) or no/mild pain (dark blue).

**Table 1:** Patients clinical and demographic information.

| Patient | Age | Sexe | Handedness | IMP<br>Lat | EZ | N°<br>contacts |
| --- | --- | --- | --- | --- | --- | --- |
| P1 | 41 | M | R | R | T (R) | 185 |
| P2 | 48 | F | R | L | T (L) | 159 |
| P3 | 24 | F | R | B | T (R) | 184 |
| P4 | 35 | F | R | L | F (L) | 183 |
| P5 | 24 | F | L | R | T & P (R) | 186 |
| P6 | 28 | M | R | L | T (L) | 185 |
| P7 | 36 | F | R | L | T (L) | 186 |
| P8 | 22 | F | 1 | R | T (R) | 178 |
| P9 | 28 | M | .67 | B | T (R) | 206 |
| P10 | 26 | M | .82 | R | F (R) | 181 |
| P11 | 32 | M | .45 | L | T (L) | 230 |
| P12 | 35 | M | .67 | R | F (R) | 204 |
| P13 | 27 | M | .09 | R | F (R) | 201 |
| P14 | 45 | M | .82 | R | P (R) | 161 |
| P15<br>(1<sup>st</sup>) | 26 | F | 1 | B | ND | 233 |
| P15<br>(2<sup>nd</sup>) | 26 | F | 1 | R | Oper. ® | 188 |
| P16 | 31 | H | .64 | R | T (R) | 153 |

*Note: the table including the number of patient (P), the age at the recording date, the sexe, the manual lateralization (Handedness, Right -R- or left -L- for the half of the patients and the manual lateralization score as evaluated with the Edinburgh inventory). Implantation lateralization (IMP Lat, right -R-, left -L- and Bilateral -B-). Epileptogenic zone location (EZ, including temporal -T-, parietal -P-, frontal -F-, Opercular -Oper-, no defined -ND-); N° Contacts, number of recorded cortical sites for each patient.*

### HFA response selectivity and graduation to sensation sentences in the posterior insula

The Semantic Categorization task revealed singular HFA responses to sensation sentences in posterior insular cortex in 5 of 6 patients, clearly functionally distinct from its anterior counterpart (Figure 1.C). Sensation sentences triggered an early and pronounced increase of HFA in the posterior insula (150-400 ms from word onset, Wilcoxon, FDR corrected p < .05), while no significant responses were detected for action or abstract sentences (Wilcoxon, FDR corrected p > .05). Among these five responsive patients, two exhibited a significant difference of HFA responses between sentence categories, specifically distinguishing sensation from abstract or action sentences (P8 and P16). In contrast, anterior insula did not elicit specific responses, showing instead a delayed (400-750 ms) and equal HFA increase for all sentence categories, in 13 of 14 patients). This inter-patient replicability is further corroborated by a group-level analysis, which shows a significant HFA increase in the posterior insula exclusively for sensation sentences (FDR corrected p < .01), differing significantly from the two other sentence categories (abstract and action, Figure1.C). Compelling evidence for the stability of the response pattern in posterior insula emerged from a patient (P15) whose dual recording sessions, separated by three months involving no brain tissue resection and a different electrode implantation site within the posterior insula, showed identical HFA selectivity for sensation sentences across both sessions (Figure 1.D).

To confirm the observed effects in posterior insula, we focused on the 6 responsive recordings from 5 patients and assessed a more sensible measure of this phenomena through single-trial response concentration (i.e. ΔQ4-Q1, quartile difference of ranked mean amplitude, see STAR methods) across conditions and sequential time intervals. A repeated measures ANOVA (Timing X Condition) revealed that trial concentration was also selectively modulated during sensation sentences reading. More specifically, we found no main effect of Timing (F(5, 25) = 1.939, p = .123, 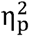 = .279), but a main effect of Condition (F(2, 10) = 5.489, p = .025, 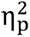 = .523) and a Timing by Condition interaction (F(10, 50) = 2.997, p = .005, 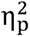 = .375). This interaction was characterized by a concentration difference between the lowest and highest quartiles, observed exclusively in the 156–297 ms interval for sensation sentences (ΔQ4-Q1 = 17.63 ± 3.69), with significant difference when compared to both abstract (ΔQ4-Q1 = -8.31 ± 4.45) and action sentences (ΔQ4-Q1 = -7.32 ± 3.89, Bonferroni corrected p < .01, Figure 1.C). Coherently, the only significant difference occurred in a time window aligned with the selective HFA response time window (to see values for all timing, see Table S2 and Figure S2). Interestingly, no such effect was observed in the anterior insula across all type of semantic categories (FDR corrected p = .42). Taken together, these findings strongly suggest that HFA responses in the posterior insula exhibit response selectivity to sensation-related sentences, characterized by a transient increase in both HFA responses and single-trial concentration. Moreover, the striking consistency enhances the reliability and validity of the posterior insula selectivity for sensation-related sentences.

This functional dissociation between the posterior and the anterior insula aligns with existing literature identifying functionally distinct regions in the human insula^18–21^. While the posterior insula has been consistently associated with pain and interoceptive sensations processing^19–21,31^, the anterior insula has proposed as a central hub for the multimodal integration^18^. Interestingly, our results are in strong agreement with previous research reporting posterior insula implication during actual pain perception^18,19,21^. What is particularly compelling is the near-perfect spatial overlap of the posterior insula activations in our study with those reported in the literature for pain perception (mean: ±X = 36, Y = -20, Z = 11; ±X = 34, Y = -20, Z = 14^18^). This remarkable spatial concordance bolsters the strength and consistency of our findings, demonstrating that pain information, whether caused physically or elicited by a word, could involve the activity of the same neuronal assemblies. To note that HFA response selectivity was detected in 20% (n = 7 of 35) and 32% (n = 35 of 109) of the posterior and anterior insula sites, respectively, indicating that HFA responses during words reading concentrate within specific neuronal populations, rather than being uniformly distributed across the insula. Although the effects of touch-related words on tactile perception^32^ and pain-related words on neural activity using fMRI^15–17^ have been previously investigated, our study provides the first direct intracranial evidence of posterior insula implication during the processing of sensation-related sentences.

Finally, we explored whether the posterior insula’s HFA selectivity could be influenced by the quality of perceived pain intensity evoked by the different sensation sentences. Therefore, patients rated each sensation-related sentence on a scale from 1 to 5 (1 = no pain, 5 = very intense pain). In 3 of 5 patients, sentences describing moderate to very intense pain elicited significantly stronger HFA modulations in the posterior insula compared to the baseline, while sentences associated with mild or no pain did not (Wilcoxon, HDR corrected p < .05). These signals thus also convey information about the subjectively perceived intensity of the sensation-related sentences. Again, the anterior insula did not show such response feature, demonstrating more generalized activity for both very intense and mild pain sentences, emphasizing the functional dissociation between the two subregions (Figure 1.E). Interestingly, these results align with prior findings reporting an intriguing correlation between the amount of posterior insula activation and the pain intensity subjectively rated by individuals^31^.

### HFA response selectivity to action sentences inside motor cortices

Figure 2.A shows evidence that HFA in the primary motor cortex (M1), located within the precentral gyrus, frequently increases around 150-400 ms when patients read sentences describing hand movements – i.e., actions-related sentences– (in 5 of 6 patients, 37% responsive electrode contacts, n = 7 of 19). Four of these patients also underwent motor localizer task, which involved actual hand, foot and face movements. This provided an opportunity to directly compare the HFA responses during real hand movements and during hand action-related sentences. In three of these four patients, we observed that HFA was significantly different from baseline activity, during the real action performance and through action-related sentences reading. These overlapping activations within M1 suggests that reading action sentences can trigger neural processes in the motor cortex similar to those activated during actual physical movement, reinforcing the idea of embodied cognition, where action language comprehension is closely tied to sensorimotor systems^4,7,12,13^. Interestingly, while the main HFA increase in the motor cortex was expectedly associated with action sentences, we also observed that abstract sentences could trigger an HFA increase in some cases (in 2 out of 6 patients). For example, one patient displayed a delayed HFA increase when reading abstract sentences, with activity peaking between 500-600 ms, compared to the 300-400 ms window for action sentences. This delayed response might reflect that motor areas may be recruited in different time intervals when processing abstract concepts, hinting at underlying complex mechanisms even for seemingly non-motor-related words. This pattern of results is in line with the view that motor cortex is not only engaged during direct physical actions but could also plays a crucial role in higher-order cognitive representations of those actions^29,30^, even through language. Moreover, the occasional engagement of M1 during abstract sentences processing re-opens exciting questions about the broader role of motor systems in the comprehension of abstract concepts^33–36^.

**Figure 2.**
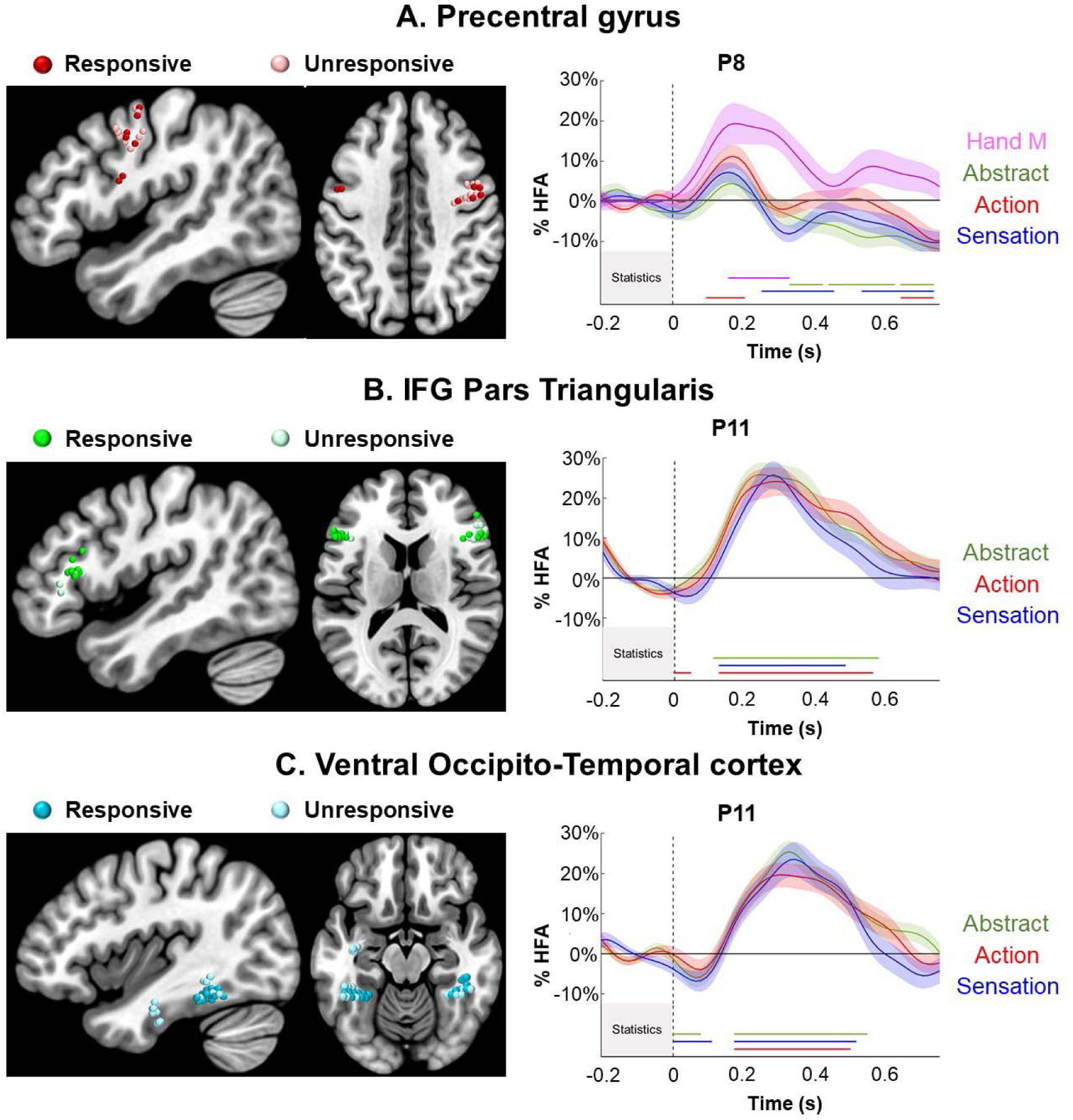
Electrode contacts localization and HFA activity in the functional control regions**: A.** the precentral gyrus (motor cortex, in red), **B.** the inferior frontal gyrus (pars triangularis in green) and **C.** the ventral occipito-temporal cortex (in cyan). Dark and light shades indicate responsives and unresponsives electrode contacts, respectively. For each functional region, the plots show %HFA [50–150 Hz] averaged modulations across trials for Abstract (green), Action (red) and Sensation (blue) sentences. Specifically for the precentral gyrus, activity from a motor localizer was also present (Hand Movement, Hand M in pink), Amplitude is expressed as a percentage relative to the average amplitude measured across the entire recording session, with shaded areas representing the Standard Error of the Mean (SEM). Colored lines below the HFA plot indicate statistically significant difference between the experimental conditions and the baseline, whereas the black lines indicate significant difference between experimental conditions.

### Generalized HFA responses to all words in language functional regions

Finally, Figure 2.B and 2.C illustrate generalized HFA in language functional regions: the inferior frontal gyrus at the pars triangularis (IFG Tri), associated with language and more specifically related with words semantic processing^24,25^, and the ventral occipito-temporal cortex (VOTC), involved in visual word form recognition^26–28^. HFA increased significantly for all sentence categories within the two language regions. These HFA patterns are consistently observed, with 8 of 9 in the IFG Tri (47% responsive electrode contacts, n = 16 of 34) and 6 of 7 patients in the VOTC (55% responsive electrode contacts, n = 23 of 42). These findings align with existing literature, which identifies these regions as core components of language networks, as evidenced by neuroimaging studies^26,27,37–39^, as well as direct electrical stimulation and intracranial recordings^40^.

## Conclusion

Taken together, these results suggest that the human posterior insula is not only involved in the perception of actual physically-induced sensation or pain but is also selectively engaged when processing language describing bodily sensations, such as pain (as supported by fMRI evidence^15–17^). This selective engagement of the posterior insula was consistently supported through various analyses demonstrating selective positive modulation of high frequency activity for sensation-related sentences. Furthermore, the inter- and intra-patient replicability provides compelling insights into the functional dissociation between the anterior and posterior insula. This selective neural engagement occurring during an early time window for sensations-related sentences, suggests that the posterior insula contributes to lexico-semantic access. Importantly, the generalized responses observed in linguistic and perceptual regions, alongside specific responses in motor areas for action sentences, underscore the distinct and specialized role of the posterior insula in sensation words processing. The current study offers the first direct intracranial evidence supporting embodiment in language processing, particularly during silent reading of words associated with sensation or painful experiences. Our results open new avenues for understanding how the brain integrates sensory and linguistic information, offering important insights into the neural mechanisms underlying language embodiment. Nonetheless, although these findings are groundbreaking, the limited number of intracranial recordings in the posterior insula warrants further replication in future studies.

## STAR□Methods

### Key resources table and resources availability

| Software and algorithms | SOURCE | IDENTIFIER |
| --- | --- | --- |
| MATLAB | https://fr.mathworks.com/products/matlab.html | 2023a |
| Fieldtrip Toolbox | https://www.fieldtriptoolbox.org/ | v.20230913 |
| SPM12 Toolbox | https://www.fil.ion.ucl.ac.uk/spm/software/spm12/ | N/A |
| MRICroGL | https://www.nitrc.org/projects/mricrogl | v.1.2.20220720 |
| Eprime | https://pstnet.com/products/e-prime/ | 3.0 |
| Code | On request | N/A |

For further requests, contact should be directed towards Marcela Perrone-Bertolotti. This study did not generate new reagents or materials. The codes used to analyze and graphically present the results are available on request.

### Participants

Intracranial recordings were obtained from 16 neurosurgical patients with drug-resistant epilepsy (8 women, mean age: 31, range 22-48) at the Epilepsy Department of the Grenoble Neurological Hospital (CHUGA, Grenoble, France). All patients were stereotactically implanted with multilead EEG depth electrodes to localize seizure sites for subsequent clinical treatment. All patients were French native speakers, and provided an oral informed consent before their inclusion in the present study and signed a non-opposition file in the context of the MAPCOG sEEG study (IdRCB: 2017-A03248-45) as approved by the National French Science Ethical Committee and in accordance with the principles of the Declaration of Helsinki. The Edinburgh inventory^41^ was used to assess the laterality of participants (see Table 1 for details). All electrode contact data exhibiting pathological waveforms were discarded from the present study. This was achieved in collaboration with the medical staff, using visual inspection of the recordings and systematically excluding data from any electrode site that was a posteriori found to be located within the seizure onset zone.

### Stimuli and behavioral task

Participants were instructed to perform a Semantic Categorization Task in an oddball paradigm with written sentences presentations. The written minimal sentences used in the study were divided into three different semantic categories, determined by the verb used: abstract, action, and sensation-related verbs. More specifically, we constructed a 90 French sentences list: 30 items for the abstract condition (e.g., “*I admire*”), 30 items for the action condition -that were specifically related to hand actions- (e.g., “*I squeeze*”), and 30 items for the sensation condition -including verbs describing different bodily and interoceptive feeling (e.g., “*I suffer*” or “*he pinches me*”). Psycholinguistic variables (written frequency, number of letters, number of syllables, and spelling neighbors) were controlled using the Lexique.org database^42^ and statistical analyses showed no difference between the three semantic categories (Table S3).

During the semantic task, participants were presented with written sentences in block sessions. They were instructed to press a keyboard key when the written sentences presented on the screen were from a different semantic category than the one initially indicated at the beginning of each block session. Indeed, sentences from the same semantic category were presented in each block, thus constituting three different types of blocks: four Abstract blocks, four Action blocks and four Sensation blocks. A total of 12 blocks, including 22 trials were presented with a total of 264 trials. Each item was presented two times during the experiment. In each block, volunteers could encounter 15 valid sentences (e.g., action sentences inside action blocks) and 7 distractor trials (e.g., an abstract sentences appearing in the action condition, and inversely). Participants were instructed to detect “distractors” (i.e., sentence that did not correspond to the semantic category of the announced block) by press a key on the keyboard (“c”). Sentences were presented in black on a white background, and in the first-person present tense. However, some sensation-related sentences were presented in third-person to evoke sensations generated by others, and prevent the description of actions performed by patients (e.g., “*he pinches me*”). All stimuli order was randomized and counterbalanced. Each trial started with a fixation cross (1500ms), followed by a pronoun (500ms), and then the target verb (1500ms, Figure 1.A).

The experimental session took place in patients’ hospital rooms. Stimuli were presented to the participant on a 15-inch computer screen at 60 cm viewing distance. Reading task was delivered via the E-prime 3.0 software (Psychology Software Tools, Pittsburgh, PA), which managed the generation of triggers on SEEG signal.

### Electrode implantation

Fourteen to nineteen semi-rigid multi-lead electrodes were stereotactically implanted in each patient (SEEG^43^). The SEEG electrodes had a diameter of .8 mm and, depending on the target structure, consisted of 8-18 contact leads (2 mm wide and 1.5 mm apart^44^, DIXI Medical Instruments). The electrode locations selection was entirely based on clinical considerations, without any reference to the current experimental protocol. This approach introduced a significant inter-individual variability in cortical sampling across participants. Each electrode contact was identified using coregistration of pre-MRI and post-CT scan images. MNI coordinates were computed using the SPM toolbox (SPM12, Wellcome Department of Imaging Neuroscience, University College London) and the position were visually displayed using a custom script (Figure S1).

### Intracranial recordings

Intracranial EEG recordings were performed using a video-SEEG monitoring system (Micromed), which allowed the simultaneous data recording from 256 depth EEG recording sites. For all patients, data were sampled at 512 Hz with an in-line bandpass filtered from 0.1 to 200 Hz. At the time of acquisition, data are recorded using a reference electrode located in white matter, and each electrode trace is subsequently re-referenced with respect to its direct neighbor (bipolar derivations). This bipolar montage has a number of advantages over common referencing. It helps eliminate signal artifacts common to adjacent electrode contacts (such as the 50 Hz mains artifact or distant physiological artifacts) and achieves a high local specificity by canceling out effects of distant sources that spread equally to both adjacent sites through volume conduction. The spatial resolution achieved by the bipolar SEEG is on the order of 3 mm^43–45^.

### Data preprocessing and statistical analysis

#### Preprocessing

SEEG data preprocessing was conducted via Matlab (The MathWorks, Natick, Massachusetts, USA) and the Fieldtrip toolbox^46^. Data preprocessing consisted of a series of steps previously used by our team^47–49^. As previously mentioned, we first applied a bipolar set-up by re-referencing each recording site to its direct neighbor. Then, we converted raw data into HFA time-series. Indeed, as proposed by Lachaux et al. (2012), HFA is a reliable physiological marker of neuronal population activity and an index of cognitive processing^23^. To do this, we performed the following procedure^47,48,50,51^: (1) Continuous SEEG signals were first bandpass-filtered into consecutive series of 10 Hz frequency bands using a zero-phase forward and reverse filter (e.g., 10 frequency bands beginning with 50–60 Hz and ending with 140–150 Hz). (2) Then, we computed the envelope using standard Hilbert transform for each bandpass-filtered signal^52^. (3) Next, the resulting envelope was downsampled to 64 Hz (i.e., one sample every 15.625 ms). (4) Afterward, for each band this envelope signal (i.e., time-varying amplitude) was divided by its means across the entire recording session and multiplied by 100. This procedure yields instantaneous envelope values expressed in percentage (%) of the mean. (5) The envelope signals (expressed in %) for each successive frequency bands were averaged together to provide one single time series (HFA) across the entire experimentation. (6) Finally, the HFA time-series was epoched into data segments centered on each stimulus (0 to 750 ms after the verb presentation onset) and baseline corrected (200 ms preceding the beginning of verb presentation).

#### Electrode contacts selection

Electrode contacts located in the posterior and anterior insula, the precentral gyrus (primary motor cortex), the inferior frontal gyrus (pars triangularis), and the fusiform gyrus were selected for analysis. The anatomical and functional locations of these electrodes were confirmed using the Julich^53^ and the AICHA^54^ brain atlases, respectively.

#### Statistical analysis

To ensure that patients performed correctly the semantic decision task, we analyzed the percentage of correct responses. We excluded filler trials from the analysis, and we have selected only the target trials for which participants answered accurately to the semantic decision, to prevent movement and motor contamination during words reading. For instance, within an action condition block, participants were instructed to press a key on the keyboard only when the presented sentence did not correspond to an action condition. However, if a participant made an error and responded to an action sentence, the resulting HFA signal from that otherwise valid trial could be confounded by the motor response. To address this issue, we systematically excluded both filler trials and target trials associated with incorrect responses. Each patient performed correctly the task, with a percentage of correct responses that exceeded the chance level (50%) for each of the experimental conditions (abstract: mean = 83.53%, SEM = 4.23%, action: mean = 84.02%, SEM = 3.72%; sensation: mean = 75.73%, SEM = 3.50%). These percentage of correct responses indicate that patients successfully performed the semantic task.

Statistical analyses were conducted with Matlab (The MathWorks, Natick, Massachusetts, USA) for the HFA time series (50-150Hz) as previously described. Comparisons were performed separately for each recording sites. First, we used paired sample Wilcoxon signed rank tests. This analysis allowed the investigation of the precise onset time and duration of the HFA increases or decreases for each experimental condition (i.e., during abstract, action and sensation sentences) compare to baseline activity. Second, to compare the activity between the experimental conditions, we used paired sample Wilcoxon signed rank tests. Moreover, to examine whether posterior insula HFA responses were influenced by pain intensity evoked by sensation sentences, Patients rated the pain intensity elicited by each sensation sentence on a scale from 1 to 5 (1 = no pain, 5 = very intense pain). Based on these ratings, the items were divided into two groups: PainMax (sentences describing moderate to very intense pain) and PainMin (sentences evoking no to mild pain). As previously described, paired sample Wilcoxon signed rank tests were used to compare each condition against baseline activity, and to examine differences between experimental conditions. All analysis was followed by a false discovery rate (FDR) correction across all time samples^55^.

We further analyzed the distribution of trials exhibiting the lowest and highest HFA amplitudes within each semantic condition to gain insights into response variability. The procedure consisted of the following steps: (1) All trials were ranked in ascending order based on their HFA amplitudes and divided into quartiles (Q1 to Q4), with Q1 and Q4 corresponding to the lowest and highest HFA responses, respectively. (2) The percentage of trials in each quartile was computed for each condition (abstract, action and sensation sentences) relative to the total number of trials in that condition 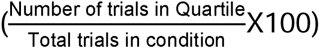. (3) To quantify the disparity in trial concentration between the lowest and highest HFA amplitudes, the difference between Q4 and Q1 (ΔQ4-Q1) was calculated for each condition. (4) This analysis was performed across multiple time windows: baseline (-156 ms to -15.6 ms), as well as post-stimulus intervals of 0–141 ms, 156–297 ms, 313–453 ms, and 469–609 ms, and 625–750 ms. (5) Since normality and sphericity were not violated (Shapiro-Wilk and Mauchly tests, respectively), a parametric repeated measures analysis of variance (ANOVA) was conducted with Timing (six time windows) and Condition (abstract, action, and sensation sentences) as factors. Post-hoc pairwise comparisons with Bonferroni corrections were applied to identify differences between conditions. The data are presented as mean values (± SEM).

## Supplementary materials

**Figure S1.**
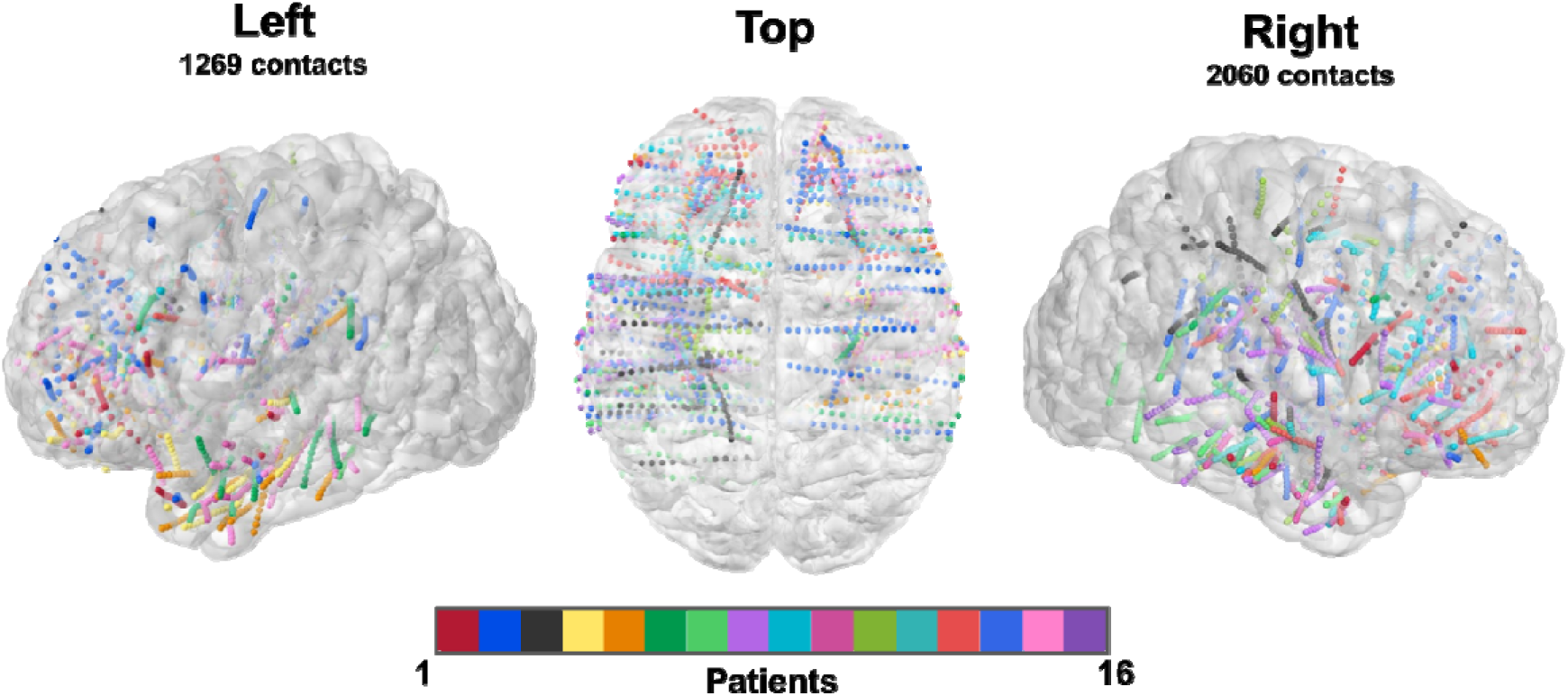
Anatomical location of all cortical sites recorded (3,329) during stereo-electroencephalographic (SEEG in 16 patients).

**Figure S2.**
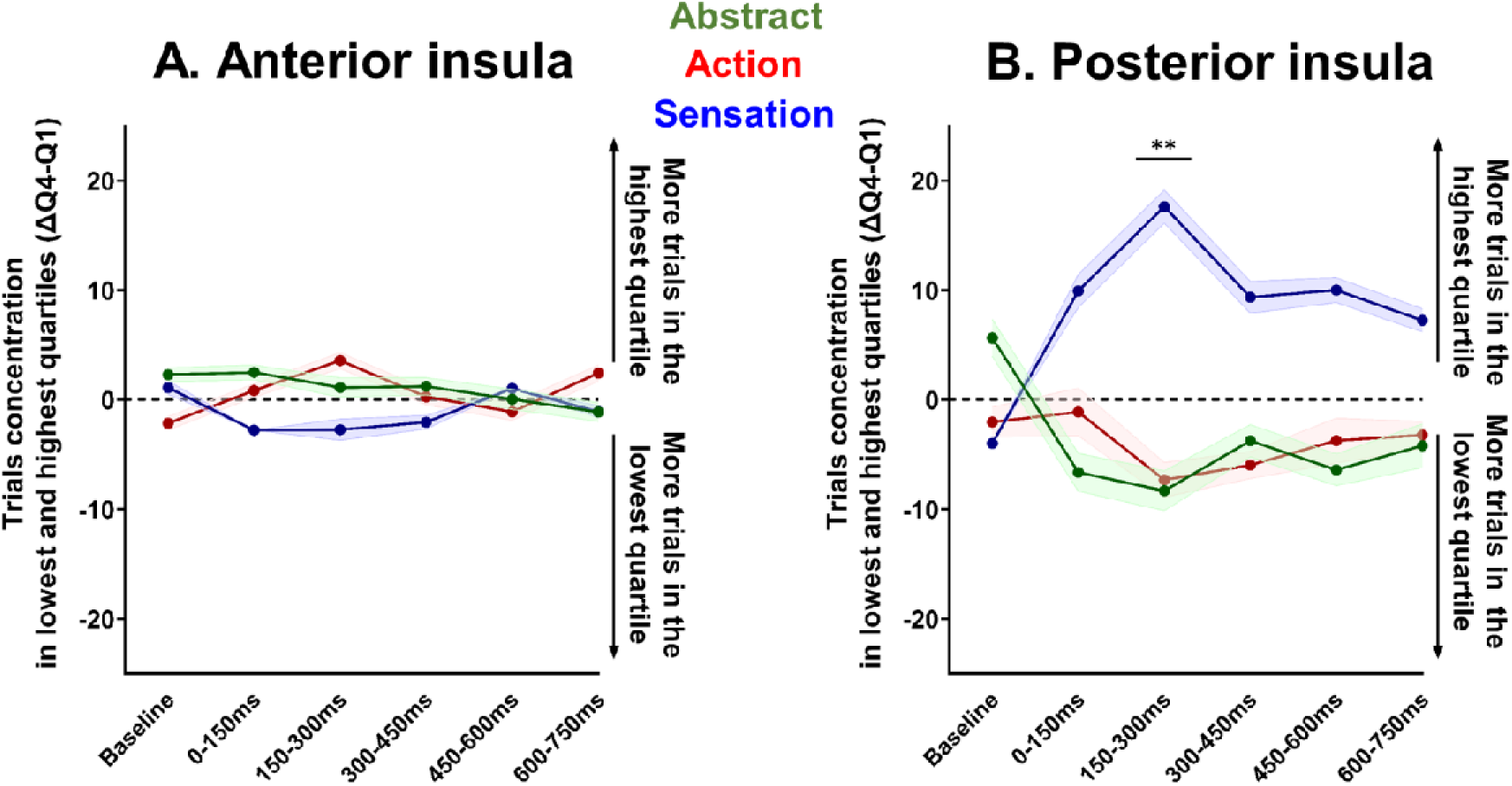
Trials concentration differences between Q1 and Q4 (ΔQ4-Q1) for each condition and each timing. ANOVA revealed a significant Timing by Condition interaction, marked by a concentration difference at 156 – 297 ms between sensation sentences and abstract or action ones.

**Table S1:** Semantic categorization task performances for each patient and each sentence condition (% of correct responses for analysis).

| Patient | Abstract | Action | Sensation |
| --- | --- | --- | --- |
| P1 | 80.00 | 63.33 | 40.00 |
| P2 | 45.00 | 53.33 | 71.67 |
| P3 | 90.00 | 98.33 | 88.33 |
| P4 | 51.67 | 93.33 | 91.67 |
| P5 | 56.67 | 81.67 | 95.00 |
| P6 | 81.67 | 93.33 | 53.33 |
| P7 | 68.33 | 65.00 | 73.33 |
| P8 | 78.33 | 73.33 | 66.67 |
| P9 | 91.67 | 93.33 | 83.33 |
| P10 | 91.67 | 100.00 | 98.33 |
| P11 | 73.33 | 86.67 | 88.33 |
| P12 | 83.33 | 96.67 | 98.33 |
| P13 | 83.33 | 96.67 | 100.00 |
| P14 | 83.33 | 61.67 | 80.00 |
| P15 (1<sup>st</sup>) | 61.00 | 78.33 | 96.67 |
| P15 (2<sup>nd</sup>) | 90.55 | 98.33 | 98.33 |
| P16 | 77.51 | 95.00 | 96.67 |
| Mean | 75.73 | 84.02 | 83.53 |
| ±SEM | 4.23% | 3.72% | 3.50% |

**Table S2:**
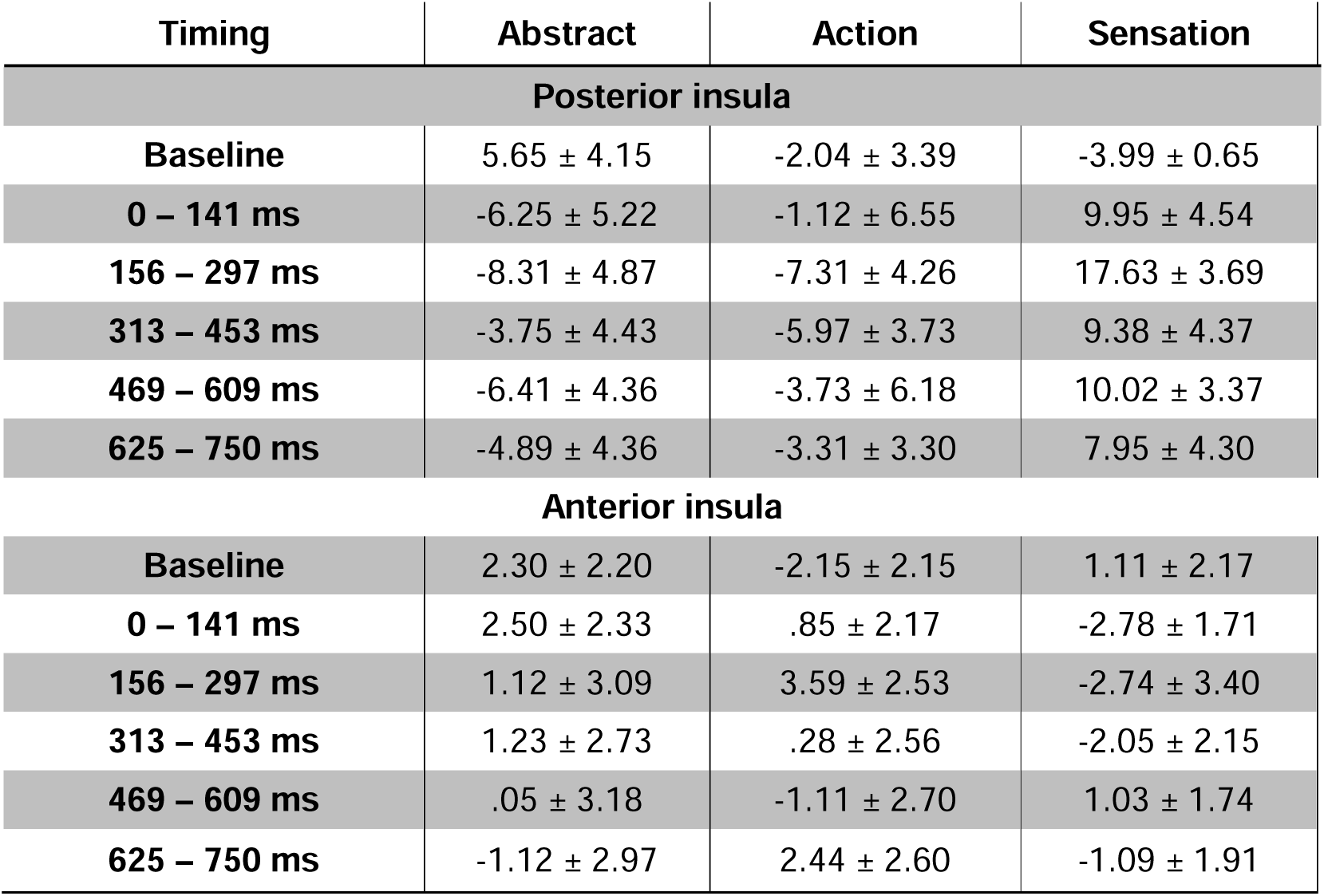
Trials concentration differences in Q1 and Q4 (ΔQ4-Q1) for abstract, action and sensation sentences at each timing.

| Timing | Abstract | Action | Sensation |
| --- | --- | --- | --- |
| **Posterior insula** |  |  |  |
| Baseline | 5.65 ± 4.15 | -2.04 ± 3.39 | -3.99 ± 0.65 |
| 0 – 141 ms | -6.25 ± 5.22 | -1.12 ± 6.55 | 9.95 ± 4.54 |
| 156 – 297 ms | -8.31 ± 4.87 | -7.31 ± 4.26 | 17.63 ± 3.69 |
| 313 – 453 ms | -3.75 ± 4.43 | -5.97 ± 3.73 | 9.38 ± 4.37 |
| 469 – 609 ms | -6.41 ± 4.36 | -3.73 ± 6.18 | 10.02 ± 3.37 |
| 625 – 750 ms | -4.89 ± 4.36 | -3.31 ± 3.30 | 7.95 ± 4.30 |
| **Anterior insula** |  |  |  |
| Baseline | 2.30 ± 2.20 | -2.15 ± 2.15 | 1.11 ± 2.17 |
| 0 – 141 ms | 2.50 ± 2.33 | .85 ± 2.17 | -2.78 ± 1.71 |
| 156 – 297 ms | 1.12 ± 3.09 | 3.59 ± 2.53 | -2.74 ± 3.40 |
| 313 – 453 ms | 1.23 ± 2.73 | .28 ± 2.56 | -2.05 ± 2.15 |
| 469 – 609 ms | .05 ± 3.18 | -1.11 ± 2.70 | 1.03 ± 1.74 |
| 625 – 750 ms | -1.12 ± 2.97 | 2.44 ± 2.60 | -1.09 ± 1.91 |

**Table S3:** T-test results concerning the linguistic and psycholinguistic characteristics of stimuli.

| **Factors** | **Abstract** | **Action** | **Sensation** | **ANOVAs** |
| --- | --- | --- | --- | --- |
| **Written frequency** | 18.78±20.96 | 24.40±63.70 | 20.27±27.27 | P= .87 ($\eta_p^2$ =.004) |
| **Number of characters** | 7.30±1.34 | 7.10±1.16 | 7.00±1.46 | P= .68 ($\eta_p^2$ =.013) |
| **Number of syllables** | 2.63±0.61 | 2.33±0.66 | 2.27±0.64 | P= .07 ($\eta_p^2$ =.088) |
| **Spelling neighbors** | 2.53±1.36 | 3.57±2.18 | 3.67±1.97 | P= .06 ($\eta_p^2$ =.093) |

